# *Leishmania infantum*-exploited Nrf2 transcription factor as a virulence process to escape macrophage-driven ferroptosis-like leishmanicidal process

**DOI:** 10.1101/2023.09.15.557964

**Authors:** Clément Blot, Kimberley Coulson, Marie Salon, Margot Tertrais, Rémi Planès, Karin Santoni, Hélène Authier, Godefroy Jacquemin, Mouna Rahabi, Mélissa Parny, Isabelle Raymond Letron, Etienne Meunier, Lise Lefèvre, Agnès Coste

## Abstract

Macrophages are the main effector cells during *Leishmania* infection. They contribute to the detection and elimination of *Leishmania spp.* and may also promote parasite resilience. Here, we report that the activation of the transcription factor Nrf2 in macrophages plays a pivotal role in the progression of *Leishmania infantum* infection by controlling inflammation and redox balance of macrophages. We also highlight the involvement of NOX2/ROS axis in the early Nrf2 activation and subsequently of PGE2/EP2r signalling in the sustainment of Nrf2 activation upon *L. infantum* infection. Moreover, we establish macrophage-driven ferroptosis-like process as a cell death program of *L. infantum* and the protective effect of Nrf2 in macrophages against *L. infantum* death. Altogether, these results identify Nrf2 as a critical factor for the susceptibility of *Leishmania infantum* infection, highlighting Nrf2 as a promising pharmacological target for the development of new therapeutic approaches for the treatment of visceral leishmaniasis.

## Introduction

*Leishmania* species are intracellular protozoan parasites that cause multiple diseases ranging from nonlethal cutaneous leishmaniasis to severe visceral disease if untreated (Mann *et al*, 2021). *Leishmania infantum* is one of the major parasite species associated with visceral leishmaniasis (Murray *et al*, 2005). Macrophages are the major myeloid immune cells which contribute to the detection and the elimination of *Leishmania spp*. However, they are also the primary replication sites for these parasites by providing them an environment suitable to their life cycle (Kaye & Scott, 2011). Macrophages can control *Leishmania infantum* infection through the generation of reactive oxygen species (ROS) (Lefèvre *et al*, 2013), underscoring the critical role of redox balance in determining the disease’s clinical outcome. The transcriptional master regulator of cellular responses against oxidative stress, known as Nuclear factor erythroid 2-related factor 2 (Nrf2), is among the key factors that regulate the redox balance (Vomund *et al*, 2017). Indeed, Nrf2 controls the expression of a multitude of antioxidant and phase II enzyme genes (Ngo & Duennwald, 2022). Furthermore, Nrf2 activation limits inflammation by decreasing the transcription of pro-inflammatory cytokines through NFκB-dependent and - independent ways (Lin *et al*, 2008; Thimmulappa *et al*, 2006).

A wide variety of intracellular pathogens such as viruses, bacteria and protozoan parasites enhances Nrf2 activation, hence leading to immune tolerance (Bichiou *et al*, 2021; Herengt *et al*, 2021; Pang *et al*, 2022). It is well established that cutaneous and visceral forms of leishmaniasis upregulate Nrf2 pathway in macrophages in response to parasite-induced ROS production (Vivarini & Lopes, 2019a). Thus, in the early stages of parasite infection, ROS-generated activation of Nrf2 constitutes a strategy developed by the parasite to subvert exposure to oxidants and to survive in macrophages (Reverte *et al*, 2021).

The regulation of Nrf2 by *Leishmania* strains increases SOD1 and HO-1 antioxidant genes, thereby promoting parasite persistence. Indeed, the establishment of a visceral infection by *L. donovani* is dependent on the induction of HO-1 by Nrf2 (Saha *et al*, 2021; Vivarini & Lopes, 2019b). This leads to a decrease in cellular heme content, thus inhibiting the maturation of NADPH oxidase subunits which results to the suppression of ROS production and participates in parasite survival in macrophages (Saha *et al*, 2019). Although some data have linked macrophage ROS production to *Leishmania* elimination, no study has addressed cell death program involved in ROS-mediated killing of *Leishmania* (Lefèvre *et al*, 2013). Interestingly, glutathione peroxidase 4 (GPX4), which is an identified transcriptional target of Nrf2, is crucial in the regulation of iron-dependent non-apoptotic mode of cell death termed ferroptosis (Song & Long, 2020). Indeed, GPX4 prevents the accumulation of toxic lipid ROS and thereby blocks the onset of ferroptosis (Anandhan *et al*, 2020; Dodson *et al*, 2019). Interestingly, the lethal phenotype of *Trypanosoma brucei* that lacks the tryparedoxin peroxidases, distant relatives of GPX4 in higher eukaryotes (Bogacz & Krauth-Siegel), strongly suggests that ferroptosis cell death program can occur in *Leishmania spp*. parasite (Bagayoko & Meunier, 2022).

It is true that the activation of Nrf2 by *Leishmania* strains is well documented, but the precise molecular mechanisms involved in this activation are not yet fully understood (Vivarini & Lopes, 2019a). However, a previous study has proposed that *Leishmania donovani* facilitates an immunosuppressive environment in macrophages through prostaglandin E2 (PGE2) / EP2 receptor signaling (Saha *et al*, 2014). Moreover, PGE2 secretion by macrophages in response to many microbial infections, such as Kaposi’s sarcoma-associated herpes virus (KSHV) viral infection, is also reported. This PGE2 production upon viral infection leads to activation of Nrf2, resulting in a permissive microenvironment for KSHV (Gjyshi *et al*, 2014).

Based on current findings that underscore the crucial role of Nrf2 in macrophage-mediated control of *Leishmania infantum* infection, we hypothesized that the oxidative burst and PGE2 secretion by *Leishmania*-infected macrophages could activate Nrf2, thereby promoting the infection’s progression by protecting the parasite against ferroptotic cell death.

The objective of our study was to investigate the role of the Nrf2 pathway in *Leishmania infantum* infection. Specifically, we aimed to elucidate the molecular mechanism responsible for activating Nrf2 signalling in macrophages in response to *Leishmania infantum.* Pharmacological inhibition or genetic deletion of Nrf2, allowed us to identify the transcription factor Nrf2 as a major player in the progression of *Leishmania infantum* infection through its impact in the control of inflammation and redox balance of macrophages. Our findings have shed light on the crucial signalling pathways involved in the activation and sustenance of Nrf2 activation during *Leishmania infantum* infection. Specifically, we have identified the NOX2/ROS axis as significant in the early stages of infection, while the PGE2/EP2 axis plays a pivotal role in the maintenance of Nrf2 activation in the later stages. Interestingly, we also demonstrated for the first time that Nrf2 in macrophages protects *L.i*. from lipid peroxidation preventing the death of parasites by ferroptosis-like process. Finally, these pathways also operated during infection of primary human macrophages, promoting Nrf2 as potential target for treatment of visceral leishmaniasis.

## Results

### *Leishmania infantum* triggers ROS and PGE2 production leading to Nrf2 activation

We have previously shown that ROS production affects *Leishmania* survival (Lefèvre *et al*, 2013). Based on these results, we investigated mRNA level of NADPH oxidase subunits and iNOS in macrophages from C57BL6 mice in response to *L. i* infection (Fig. 1). *Cybb*, *p47*, *p67* and *Nos2* mRNA levels were upregulated in macrophages challenged with *L.i.* (Fig. 1a). In line, the production of ROS and NO by macrophages following *L.i* challenge were also strongly increased (Fig.1b-c).

**Figure 1.**
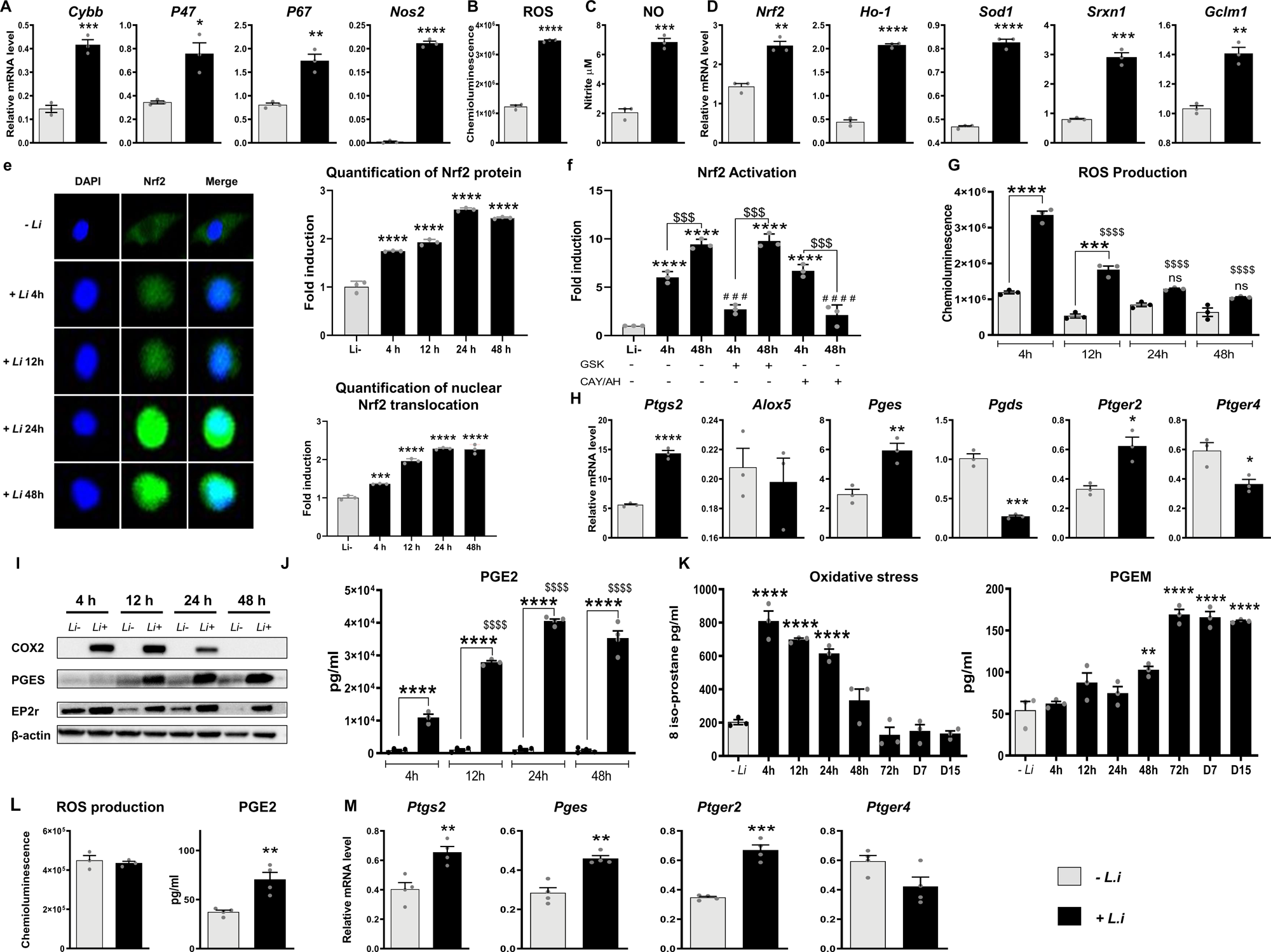
*Leishmania infantum* triggered ROS and PGE2 production leading to Nrf2 activation. A-J All analyses were performed *in vitro* on peritoneal macrophages from C57BL/6 mice in response to *L.i*. challenge. A qRT-PCR analysis of Nox2 subunits and *Nos2* genes after 4h of *L.i* challenge. B ROS production in response to *L.i* challenge. C NO production in response to *L.i* challenge. D qRT-PCR analysis of *Nrf2* and its target genes 4h post *L.i* challenge. E Confocal imaging of the Nrf2 cellular localization (green) at 4h, 12h, 24h, and 48h post-infection (nuclei stained with DAPI, blue). The top histogram represents the quantification of Nrf2 protein and the bottom histogram represents the nuclear translocation of Nrf2, both compared to uninfected macrophages (-*L.i.*). The data were represented in fold induction relative to uninfected peritoneal macrophages. F DNA-binding ELISA quantification of the ARE-nuclear binding of Nrf2 in peritoneal macrophages pre-treated with a NOX2 inhibitor (GSK2795039) or with a PGEs inhibitor (CAY10526) and an EP2 antagonist (AH6809) and infected with *L.i*, for 4h or 48h. The data were represented in fold induction relative to untreated and uninfected peritoneal macrophages. G ROS production at 4h, 12h, 24h and 48h after *L.i* challenge. H qRT-PCR analysis of Arachidonic acid metabolism enzymes genes 4h post *L.i* challenge. I Immunoblot analysis of COX2, PGEs, EP2r proteins after 4h, 12h, 24h and 48h post *L.i* challenge. J ELISA quantification of PGE2 production by peritoneal macrophages at 4h, 12h, 24h, 48h post *L.i* challenge. K-L C57BL/6 mice were infected (i.p.) with 50 × 10^6^ *L.i* for 14 days. K Urine from infected mice was collected at 4h, 12h, 24h, 48h, 72h, 7d and 14d for determination of PGE2 and 8-isoprostane levels by ELISA. L ROS and PGE2 production by peritoneal macrophages from infected mice for 14 days. M qRT-PCR analysis of Arachidonic acid metabolism genes 14 days after infection in peritoneal macrophages. Data information: Results correspond to mean ± SEM and are representative of at least three independent experiments. * p < 0.05, ** p < 0.01, *** p < 0.001 and **** p < 0.0001 compared to the uninfected macrophages (−*L.i*). ^$^ p < 0.05, ^$$^ p < 0.01, ^$$$^ p < 0.001 and ^$$$$^ p < 0.0001 compared to the infected macrophage at 4h (4h + *L.i*). ^#^ p < 0.05, ^#^ ^#^ p < 0.01, ^#^ ^#^ ^#^ p < 0.001 and ^#^ ^#^ ^#^ ^#^ p < 0.0001 compared to corresponding control (+ *L.i.* 4h untreated).

Among the factors controlling redox balance, the transcription factor Nrf2 is a major regulator of the cellular antioxidant response (Vomund *et al*, 2017). Interestingly, in macrophages from C57BL/6 mice, Nrf2 mRNA and protein levels and its target genes were upregulated during *L.i* infection (Fig. 1 d-e). This was supported by gradual increase of Nrf2 nuclear translocation from 4h to 48h post-*L.i* challenge (Fig. 1e) and by DNA-binding activity of Nrf2 at 4h and 48h post-*L.i* challenge (Fig. 1f).

It is well established that ROS are key factors of Nrf2 activation (Ngo & Duennwald, 2022). We showed that the strong induction of ROS production at 4h after *L.i* challenge was abolished at 24h (Fig. 1 g) while Nrf2 remained activated until 48h post-*L.i* challenge (Fig. 1 e-f).

Among the immunoregulatory mediators produced during *Leishmania* infection, Prostaglandin E2 (PGE2) was recently identified to facilitate parasite survival(Saha *et al*, 2014). Interestingly, we observed here in macrophages challenged with *L.i* a specific increase of mRNA and protein levels of COX-2, PGES, which are arachidonic acid metabolism enzymes involved in PGE2 production and EP2, a PGE2 receptor (Fig. 1h-i). Consistently, PGE2 production by macrophages was progressively increased from 4h post L.i challenge (Fig.1j).

To clearly establish that in the early stages of infection, Nrf2 activation in macrophages is dependent on ROS and in the late stages on PGE2 production, we evaluated the DNA-binding activity of Nrf2 in macrophages pre-treated with GSK2795039 (NOX2 inhibitor) or with CAY10526 (PGES inhibitor) associated with AH6808 (EP2 antagonist) (Fig. 1f). Interestingly, at 4h post-*L.i* challenge, only the inhibition of ROS production reduced Nrf2 translocation. Inversely, at 48h post-*L.i* challenge, only the inhibition of PGE2 pathway decreased Nrf2 activation (Fig. 1f).

These data were then supported in a murine model of *L.i*. viceral infection where we demonstrated a high concentration of urinary 8-isoprostane, which reflects the oxidative stress, in the early stages post-infection (4h-24h) and a drastic decrease of urinary 8-isoprostane from 48h post-infection and in the late stages of infection (days 7 and 15 post-infection) (Fig. 1k). Inversely, the PGE2 metabolite levels in urine increased only after 48h post-infection (Fig. 1k). This level remained elevated until day 15 after infection. The lack of ROS induction and increased PGE2 level associated to *Ptgs2*, *Pges*, and *Ptger2* mRNA overexpression in peritoneal macrophages at day 15 post-*L.i* infection (Fig. 1l-m) strengthened the contribution of ROS in early Nrf2 activation and the later involvement of PGE2 in maintaining Nrf2 activation during *L.i* infection.

### ROS- and PGE2-mediated Nrf2 activation promotes *Leishmania* proliferation

To evaluate the contribution of ROS- and PGE2-mediated Nrf2 activation in the control of *L.i.* proliferation by macrophages, we determined *L.i* proliferation in macrophages deficient for Nrf2 (Nrf2^-/-^). *L.i* proliferation was prevented in macrophages lacking Nrf2 and the inhibition of ROS production by GSK2795039 restored *L.i.* proliferation in Nrf2^-/-^ macrophages (Fig. 2a).

**Figure 2.**
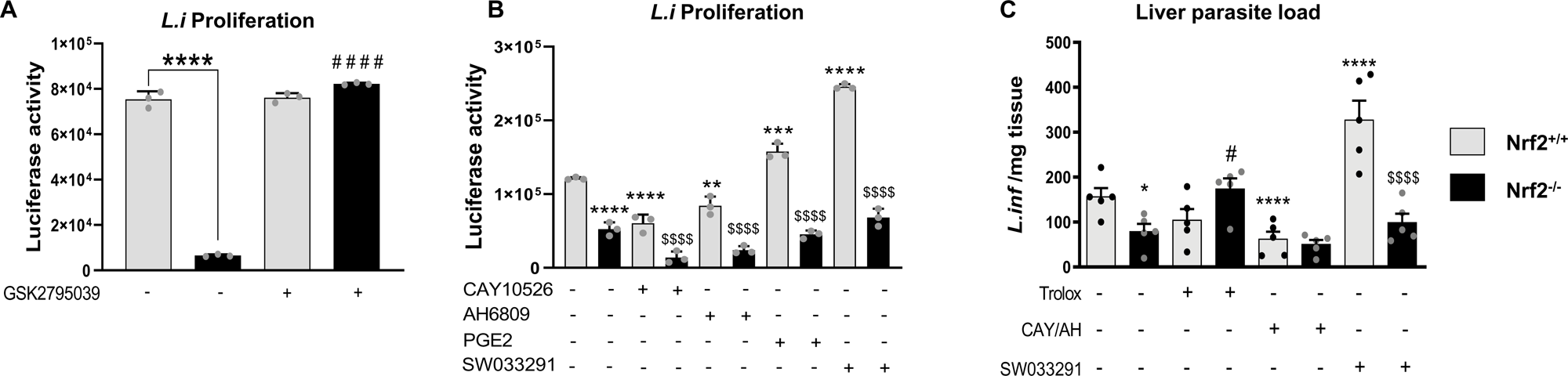
ROS and PGE2-mediated Nrf2 activation promoted *L.i* proliferation. A-B *L.i.* proliferation in peritoneal macrophages from Nrf2^+/+^ and Nrf2^-/-^ treated with a NOX2 inhibitor (GSK2795039) (A) or with PGEs inhibitor (CAY10526) associated with an EP2 antagonist (AH6809), or with 15-PGDH inhibitor (SW033291) or with PGE2 (B). Results correspond to mean ± SEM and are representative of at least three independent experiments. *p< 0.05 and ****p< 0.0001 compared to untreated macrophages from Nrf2^+/+^ mice; # # # # p<0.0001 compared to untreated macrophages from Nrf2^-/-^ mice; $$$p<0.001 and $$$$p<0.0001 compared to the respective treated macrophages from Nrf2^+/+^ mice. C Parasite load was quantified by qRT-PCR in Nrf2^+/+^ and Nrf2^-/-^ mice infected with (i.p.) with 50 × 10^6^ *L.i* for 14 days. Mice were treated with an ROS chelator (Trolox, every 2 days) or with a cocktail of PGEs inhibitor (CAY10526) and EP2 antagonist (AH6809) or with a 15-PGDH inhibitor (SW033291) (n=5). Data information: Results correspond to mean ± SEM, *p< 0.05 and ****p< 0.0001 compared to the respective untreated Nrf2^+/+^ mice; # p<0.05 compared to untreated Nrf2^-/-^ mice; $$$$p<0.0001 compared to the respective treated Nrf2^+/+^ mice.

Moreover, the inhibition of PGE2 production with CAY10526 (PGES inhibitor) and the treatment with AH6808 (EP2 receptor antagonist) decreased *L.i* proliferation only in Nrf2^+/+^ macrophages, whereas PGE2 addition increased *L.i* proliferation (Fig. 2b). We also evaluated *L.i* proliferation in macrophages treated with SW033291, a 15 Pgdh inhibitor (enzyme of PGE2 degradation), in Nrf2^+/+^ and Nrf2^-/-^ macrophages (Fig. 2b). As expected, SW033291 treatment increased *L.i.* proliferation only in Nrf2^+/+^ macrophages (Fig. 2b), demonstrating that the impact of Nrf2 on promoting *L.i.* proliferation involved specifically PGE2/EP2 axis.

Consistently with decreased *L.i* proliferation in Nrf2^-/-^ macrophages, the parasite load in liver was lower in Nrf2^-/-^ mice infected by *L.i* (Fig. 2c). Interestingly, in infected Nrf2^+/+^ and Nrf2^-/-^ mice, the treatment with Trolox (ROS chelator) increased the number of *L.i.* in the liver of Nrf2^-/-^ mice (Fig. 2c).

Conversely, the inhibition of PGE2 production by CAY10526/AH6808 reduced significantly the number of *L.i* in the liver of Nrf2^+/+^ mice (Fig. 2c). Moreover, SW033291 treatment strongly increased the parasite load in the liver of only Nrf2^+/+^ mice (Fig. 2c). Altogether, these data demonstrate that ROS and PGE2/EP2 axis mediated Nrf2 activation hence promoting *L.i.* growth.

### Nrf2 in macrophages promotes *L.i* infection through the induction of an anti-oxidant and anti-inflammatory phenotype

As the parasite loads in liver and spleen were significantly decreased in Nrf2^-/-^ mice infected by *L.i* compared to their infected wild-type littermates (Nrf2^+/+^ mice), we dissected how Nrf2 in macrophages promotes *L.i* infection in visceral murine model of leishmaniasis (Fig. 2c and 3a).

**Figure 3.**
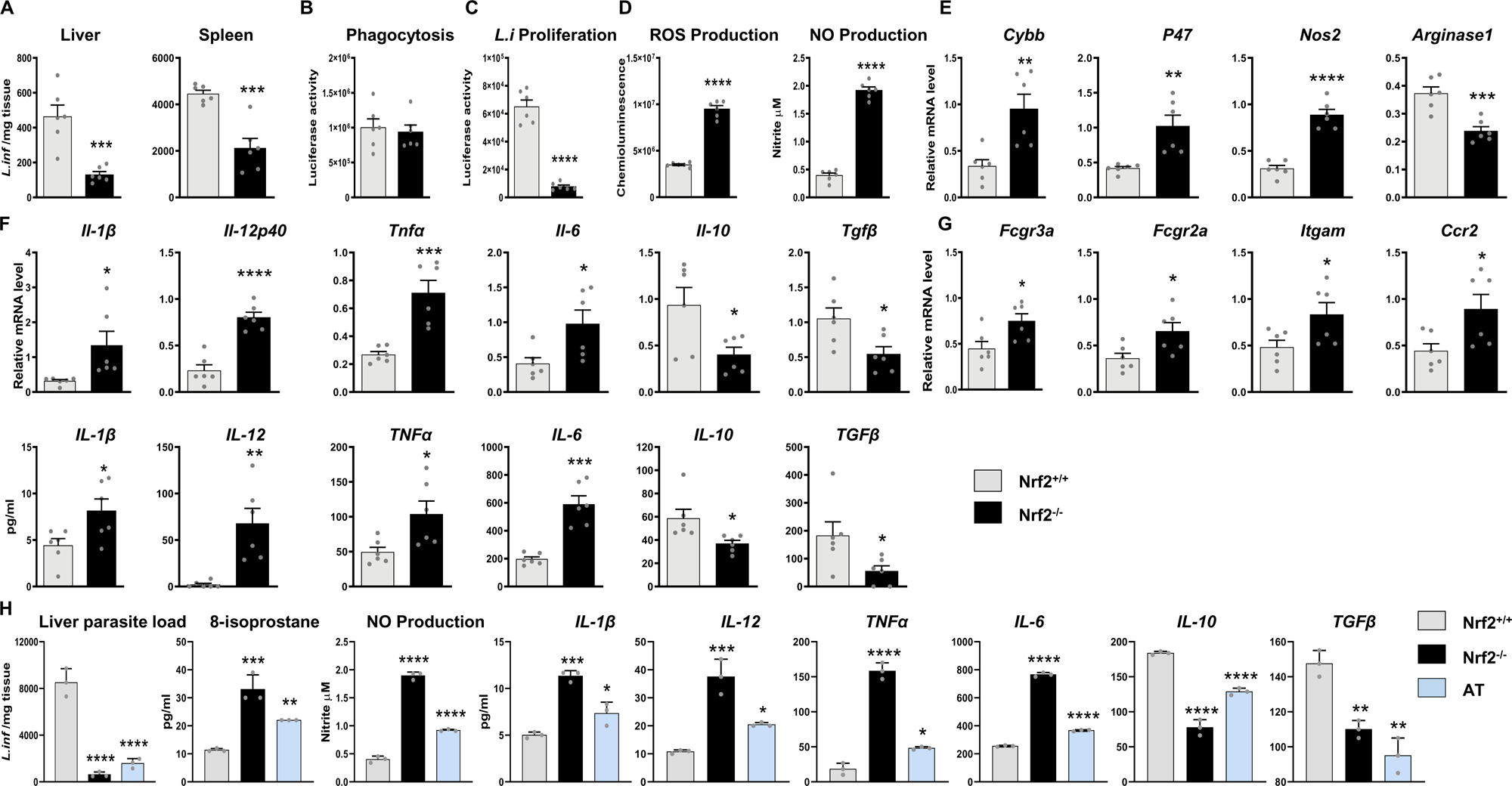
Nrf2 promotes *L. infantum* infection in macrophage *via* an anti-oxidant and anti-inflammatory phenotype. A-G Nrf2^+/+^ and Nrf2^-/-^ mice were infected (i.p.) with 50 × 10^6^ *L.i* for 14 days. A Parasite loads in liver and spleen from infected Nrf2^+/+^ and Nrf2^-/-^ mice, quantified by qRT-PCR. B-C Phagocytosis (B) and proliferation (C) of *L.i* in peritoneal macrophages from infected Nrf2^+/+^ and Nrf2^-/-^ mice. C ROS and NO productions by peritoneal macrophages from infected Nrf2^+/+^ and Nrf2^-/-^ mice. E-G qRT-PCR and ELISA analysis on peritoneal macrophages from infected Nrf2^+/+^ and Nrf2^-/-^ mice of oxidative stress markers (E), cytokines (F) and surface receptors (G). H An adoptive transfer of Nrf2^-/-^ macrophages into Nrf2^+/+^ recipient mice (AT) was performed before *L.i* infection for 14 days. Liver parasite load, 8 isoprostane and NO productions and cytokine secretion in peritoneal fluids were quantified. Data information: Results correspond to mean ± SEM and are representative of at least three independent experiments. *p<0.05, **p<0.01, *** p<0.001 and **** p< 0.0001 compared to infected mice Nrf2^+/+^.

To better characterize the role of Nrf2 in the microbicidal functions of macrophages, we evaluated *ex vivo* the ability of macrophages from Nrf2^+/+^ and Nrf2^-/-^ mice to engulf and to modulate the proliferation of *L.i* (Fig. 3b-d). Although there was no difference on phagocytosis capacity between macrophages from Nrf2^+/+^ and Nrf2^-/-^ mice, the proliferation of *L.i* was decreased in macrophages from Nrf2^-/-^ mice (Fig. 3b-c). Interestingly, ROS and NO productions of macrophages from infected Nrf2^-/-^ mice were increased in response to *L.i.* challenge compared to macrophages from infected Nrf2^+/+^ mice (Figure 3d). In line, the mRNA expression of *Cybb* and *P47* and *Nos2* were upregulated in macrophages from infected Nrf2^-/-^ mice whereas *Arg-1* mRNA level was downregulated (Fig. 3e).

Gene expression analysis and protein level of cytokines in macrophages from Nrf2^+/+^ and Nrf2^-/-^ infected mice showed a significant increase of IL-β, IL-12, TNFα, IL-6 pro-inflammatory cytokines in macrophages from Nrf2^-/-^ infected mice, which was mirrored by a decrease of IL-10 and TGFβ anti-inflammatory cytokines (Fig. 3f). Consistent with proinflammatory profile of macrophages from Nrf2^-/-^ infected mice, the *Fcgr3a*, *Fcgr2b*, *Itgam* and *Ccr2* mRNA levels were increased (Fig. 3g). These data demonstrate that Nrf2 directs macrophages towards an anti-inflammatory and anti-oxidant phenotype, promoting the progression of *L.i.* infection.

To further confirm that Nrf2 of macrophages is required for *L.i* resilience, we performed an adoptive transfer of Nrf2^-/-^ macrophages into infected Nrf2^+/+^ recipient mice (AT) (Fig. 3h). The adoptive transfer of Nrf2^-/-^ macrophages into Nrf2^+/+^ infected mice decreased parasite load in the liver while increasing microbicidal functions such as ROS, reflected by 8-isoprostane production, NO, and cytokine production in the peritoneal cavity, as observed in infected Nrf2^-/-^ mice (Fig. 3h). Thus, Nrf2 in macrophages may be the cause of systemic vulnerability to *L.i.* infection.

### Nrf2 in macrophages prevents the death of *L.i* by lipid peroxidation

To dissect how *L.i*. is eliminated in Nrf2^-/-^ mice, we evaluated whether the decrease in *L.i.* proliferation was due indirectly to macrophage death or directly to parasite mortality. We first measured *in vitro* the cell viability of Nrf2^+/+^ and Nrf2^-/-^ macrophages infected with *L.i* (Fig. 4a). *L.i* infection did not alter the macrophage viability at 24h and 48h (Fig. 4a).

**Figure 4.**
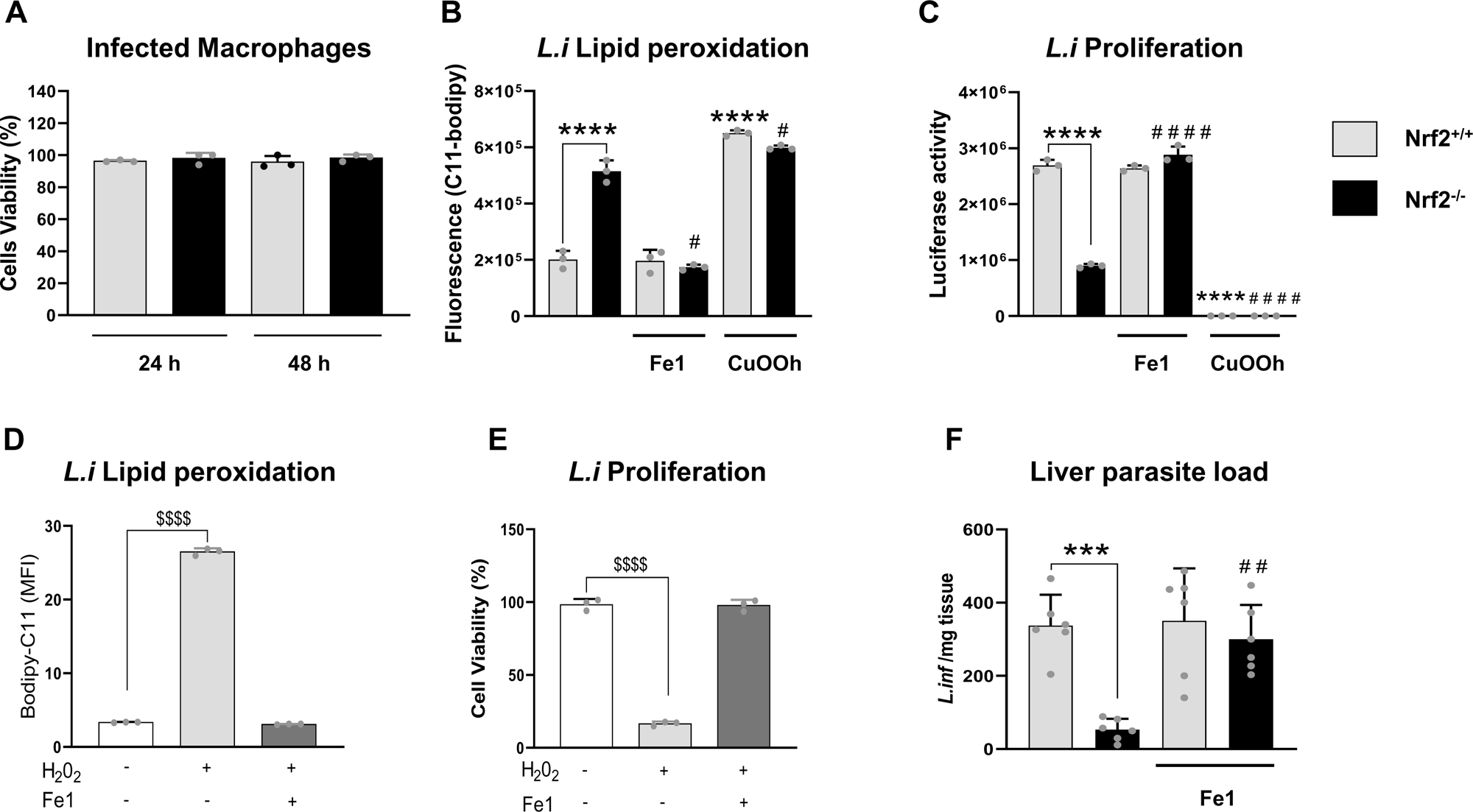
Nrf2 in macrophages prevents the death of *L.infantum* by ferroptosis. A Viability of peritoneal macrophages from Nrf2^+/+^ and Nrf2^-/-^ mice infected with *L.i.* for 24h and 48h. B-C Quantification of *L.i.* lipid peroxidation (B) and *L.i* proliferation (C) in peritoneal macrophages from Nrf2^+/+^ and Nrf2^-/-^ mice, treated or not with Ferrostatin-1 (Fe1) or Cumene hydroperoxyde (CuOOH). D Quantification of lipid peroxidation in *L.i* promastigotes incubated or not with H_2_0_2_ for 4h and treated with or without Ferrostatin-1. E Quantification *L.i* proliferation in the presence of H_2_0_2_ for 4h and with or without Ferrostatin-1. F Parasite load was quantified by qRT-PCR in Nrf2^+/+^ and Nrf2^-/-^ mice infected with (i.p.) with 50 × 10^6^ *L.i* for 14 days. Mice were treated with Ferrostatin-1 (Fe1) every 2 days. Data information: Results correspond to mean ± SEM and are representative of at least three independent experiments. **** p<0.0001 compared to untreated peritoneal macrophages from Nrf2^+/+^ mice. ^#^ p<0.05 and ^#^ ^#^ ^#^ ^#^ p<0.0001 compared to untreated peritoneal macrophages from Nrf2^-/-^ mice. ^$$$$^ <0.0001 compared to untreated *L.i* promastigotes.

It was recently described that the accumulation of oxidative stress induced peroxidation of polyunsaturated fatty acids, an essential step of ferroptosis. Moreover, glutathione peroxidase 4 (GPX4), a critical anti-ferroptotic enzyme, is a target gene of Nrf2 (Zhou *et al*, 2020). In this context, we evaluated whether *L.i.* died by ferroptosis in Nrf2*-/-* macrophages. We measured lipid peroxidation of *L.i.* using C11-BODIPY, a redox-sensitive dye, in Nrf2^+/+^ and Nrf2^-/-^ macrophages (Fig. 4b) and the number of living *Leishmania* in macrophages was estimated through their luciferase activity (Fig. 4c). We observed a higher lipid peroxidation of *L.i.* in Nrf2^-/-^ macrophages (Fig. 4b) which was associated to a decrease of *L.i.* number (Fig. 4c). Interestingly, the addition of ferrostatin-1 (Fe-1), an inhibitor of ferroptosis, prevented the *L.i.* lipid peroxidation in Nrf2^-/-^ macrophages and increased *L.i.* number in Nrf2-/- macrophages. These results were supported by an increased peroxidation of *L.i.* lipids associated with a drastic decrease in the number of *L.i.* in Nrf2^+/+^ and Nrf2^-/-^ macrophages following treatment with Cumene hydroperoxide (Cuooh), a ferroptosis inducer. These results suggested that Nrf2-induced ROS inibition protected *L.i.* from a microbicidal process highly ressembling ferroptosis.

To confirm that ROS were responsible for *L.i.* lipid peroxidation and death, we studied *in vitro* the lipid peroxidation and the viability of *L.i* in presence of H_2_O_2_, a non-ROS radical (Fig. 4d-e). In presence of H_2_O_2_, *L.i*. lipid peroxidation was strongly induced and *L.i.* viability was impaired. The addition of ferrostatin (Fe1) protected *L.i*. from H_2_O_2_-induced lipid peroxidation and improved their viability. Finally, we infected Nrf2^+/+^ and Nrf2^-/-^ mice with *Leishmania infantum*, in presence or absence of ferrostatin (Fe1). Given the very low peristance of Fe1, mice were treated each day with Fe1. We observed that the decrease in parasite load observed in Nrf2^-/-^ mice was completely abolished by Fe-1 treatment (Fig. 4f), hence suggesting that host-driven ferroptosis-like of *L. infantum* is a critical mechanism of defense against *L. infantum* in macrophages and in mice.

Altogether, those results suggested that Nrf2 in macrophages protected *L.i*. from lipid peroxidation preventing their death by a mechanism similar to ferroptosis.

### Pharmacological inhibition of Nrf2 reduces *L.i.* infection through oxidative and inflammatory profile of macrophages

To validate Nrf2 as a relevant therapeutic target for visceral leishmaniasis treatment, we used a pharmacological approach with ML-385, a Nrf2 inhibitor, in mice infected with *L.i.* (Fig. 5). Interestingly, 15 days after infection, the parasite loads in the spleen and liver of infected mice were decreased similarly when the mice were treated with ML385 or liposomal amphotericin B (AmB), a standard treatment of visceral leishmaniasis (Fig. 5a). The proliferation of *L.i* was decreased in macrophages from infected mice treated with ML385, suggesting that the pharmacological inhibition of Nrf2 promoted the microbicidal function of macrophages (Fig. 5b). Consistently, the ML385 treatment of infected mice increased ROS and NO productions of macrophages in response to *L.i.* challenge (Fig. 5c-d). In line, lipid peroxidation of *L.i* in macrophages was significantly induced by ML385 treatment (Fig. 5e). The ML385 treatment also upregulated mRNA and protein levels of IL-β, IL-12, TNFα and IL-6 pro-inflammatory cytokines and downregulated IL-10 anti-inflammatory cytokine (Fig. 5f). Thus, these data achieved with the ML385 treatment were consistent with those obtained in infected Nrf2^-/-^ mice, demonstrating the therapeutic potential of Nrf2 modulation in the treatment of visceral leishmaniasis.

**Figure 5.**
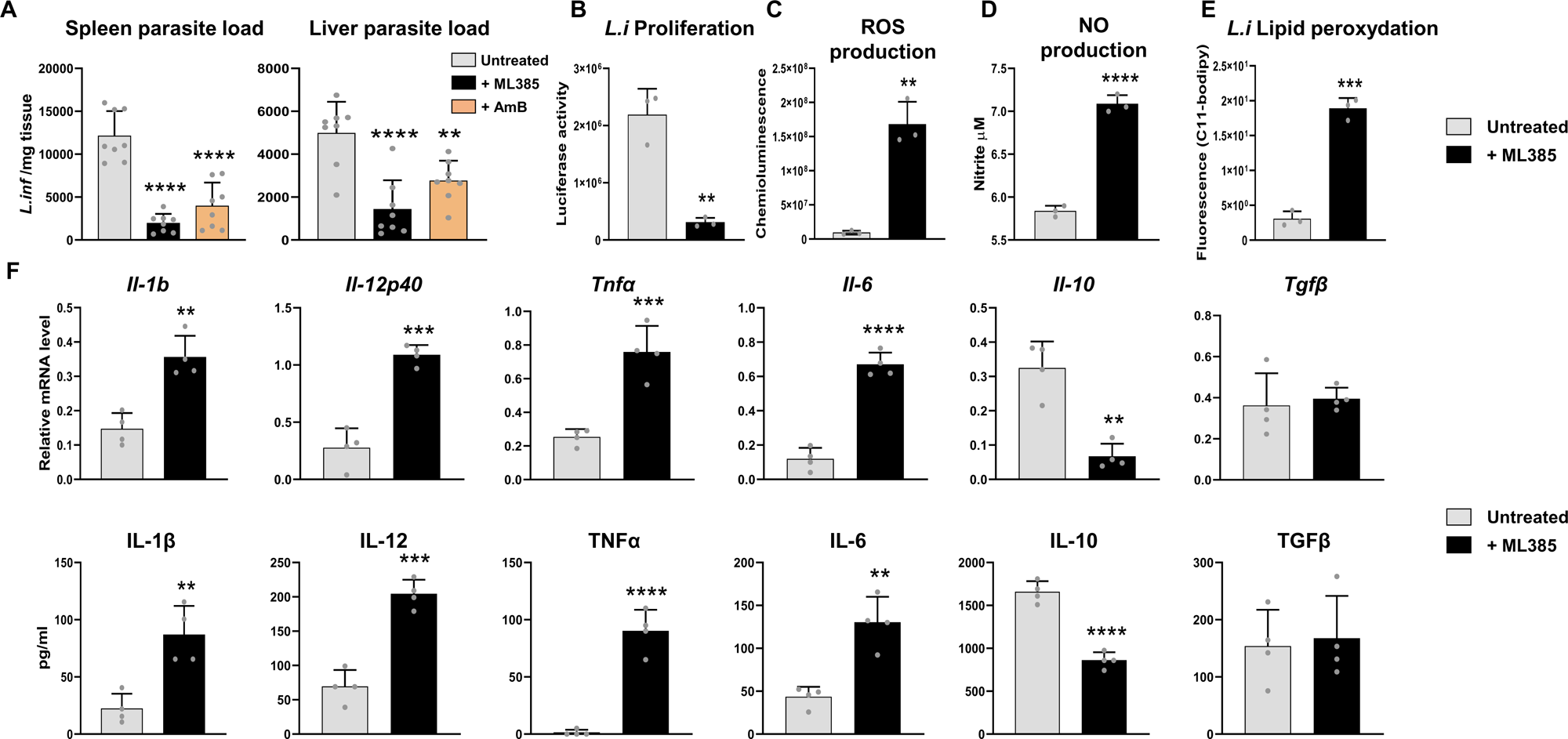
Pharmacological inhibition of Nrf2 reduces *L.infantum* infection through oxidative and inflammatory profile of macrophages. A-F C57BL/6 mice were infected (i.p.) with 50 × 10^6^ *L.i.* for 14 days and treated or not with ML385 (Nrf2 inhibitor) every 2 days or with liposomal amphotericin B (AmB) for 5 days. A Parasite loads in liver and spleen quantified by qRT-PCR. B *L.i* proliferation in peritoneal macrophages from infected mice treated or not with ML385. C ROS production by peritoneal macrophages from infected mice treated or not with ML385 in response to *L.i* challenge. D NO production by peritoneal macrophages from infected mice treated or not with ML385 in response to *L.i* challenge. E *Q*uantification of *L.i.* lipid peroxidation in peritoneal macrophages from infected mice treated or not with ML385. F qRT-PCR and ELISA analysis of cytokines on peritoneal macrophages from infected mice treated or not with ML385. Data information: Results correspond to mean ± SEM and are representative of at least three independent experiments. **p< 0.01, ***p< 0.001 and ****p<0.0001 compared to untreated mice.

### ROS/PGE2 axis-mediated Nrf2 activation enables *L.i.* proliferation in human monocyte-derived macrophages

To validate our findings from the murine model in a human context, we challenged human monocyte-derived macrophages (h-MDM) with *L.i*. (Fig. 6). The gene expression of Nrf2 and its target genes (HO-1, SRXN1, NQ01) in h-MDM were strongly increased by *L.i*. challenge (Fig. 6a). Similar to our findings in the murine model, *CYBB and P47* mRNA levels were upregulated in h-MDM challenged with *L.i.* (Fig. 6b). In line, the production of ROS by h-MDM following *L.i* challenge was increased in early stage (4h post-infection) and gradually decreased over time (Fig. 6c). Inversely, PGE2 production by h-MDM was progressively increased from 4h post *L.i.* challenge (Fig. 6e). Consistent with increased PGE2 production, *PTGS2, PGES and PTGER2* mRNA levels were upregulated following *L.i* challenge and *PGDS* and PTGER4 were downregulated (Fig. 6d). Consequently, Nrf2 nuclear translocation in h-MDM was gradually increased from 4h to 48h post-*L.i* challenge (Fig. 6f).

**Figure 6.**
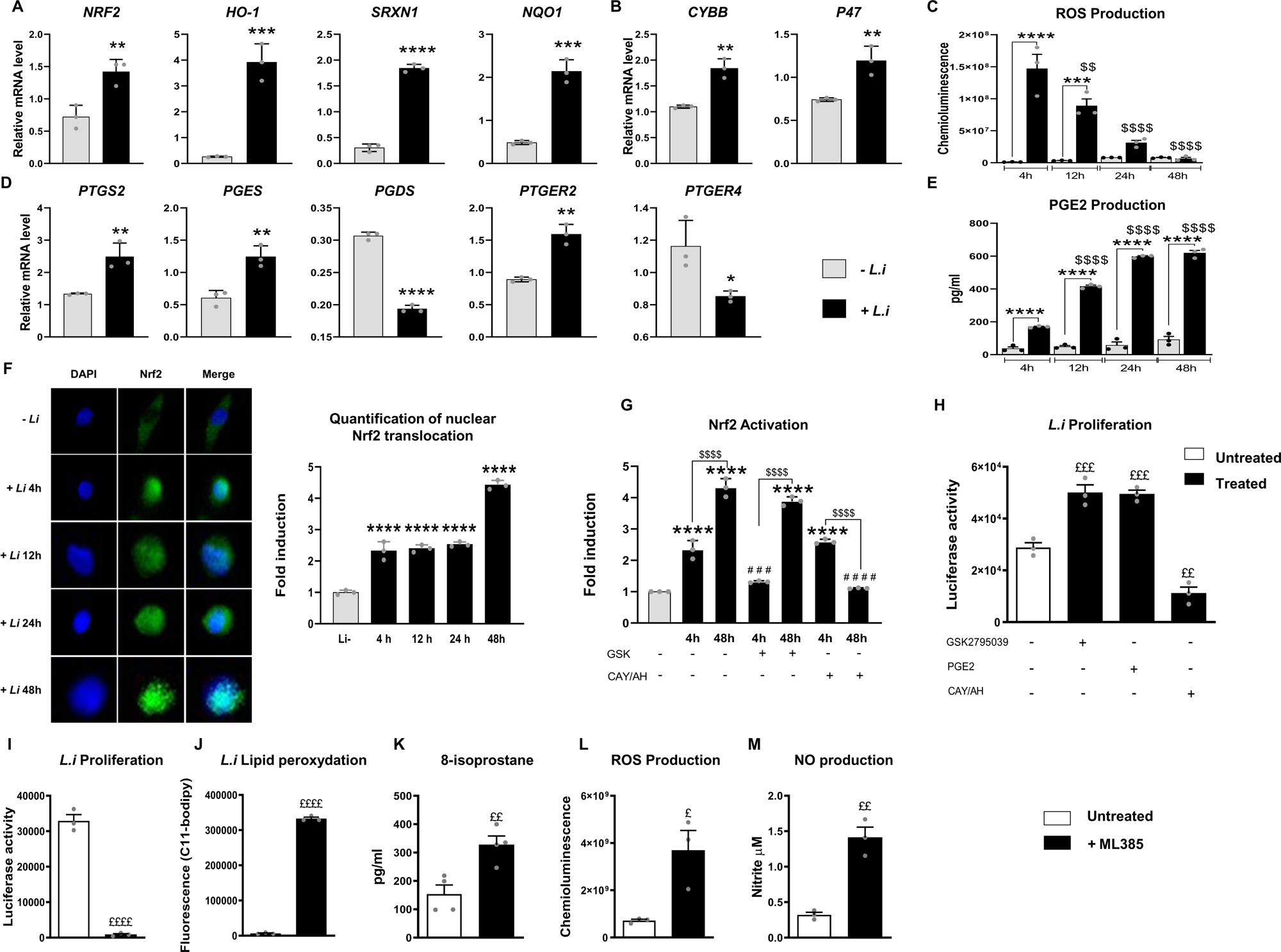
ROS/PGE2 axis-mediated Nrf2 activation enables *L.infantum* proliferation in human monocyte-derived macrophages. A-M All analyses were performed *in vitro* on human monocyte-derived macrophages (h-MDM) in response to *L.i* challenge. A-B qRT-PCR analysis of NRF2 and its target genes and NOX2 subunits 4h after *L.i* challenge. C ROS production at 4h, 12h, 24h and 48h after *L.i* challenge. D qRT-PCR analysis of Arachidonic acid metabolism genes 4h after *L.i*. challenge. E PGE2 production at 4h, 12h, 24h, 48h after *L.i* challenge. F Confocal imaging of NRF2 (green) cellular localization and DAPI-labeled nucleus (blue) at 4h, 12h, 24h, and 48h after *L.i* challenge. Quantification of nuclear translocation of NRF2 compared to uninfected h-MDM (-*L.i.*). The data were represented in fold induction relative to uninfected h-MDM. G Quantification of nuclear translocation of NRF2 by confocal imaging at 4h and 48h after *L.i* challenge in h-MDM treated or not with GSK2795039 (NOX2 inhibitor) or CAY10526 (PGEs inhibitor) and AH6809 (EP2 antagonist). The data were represented in fold induction relative to uninfected h-MDM. H *L.i* proliferation in h-MDM treated or not with GSK2795039 or CAY10526/AH6809 or with PGE2. I *L.i* proliferation in h-MDM treated or not with ML385. J Lipid peroxidation of *L.i* in h-MDMs treated or not with ML385. K 8-isoprostane quantification of *L.i* challenged h-MDMs treated or not with ML385. L ROS production by h-MDMs challenged with *L.i* and treated or not with ML385. M NO production by h-MDMs challenged with *L.i* and treated or not with ML385. Data information: Results correspond to mean ± SEM and are representative of at least three independent experiments. *p<0.05, **p<0.01, and ****p< 0.0001 compared to uninfected h-MDM (-*L.i*). ^$$$$^ p<0.0001 compared to 4h infected h-MDM (4h + *L.i*.). ^#^ ^#^ ^#^p< 0.001 and ^#^ ^#^ ^#^ ^#^p< 0.0001 compared to untreated infected h-MDM at 4h. ^£^p<0.05, ^£^ ^£^ p<0.01, ^£^ ^£^ ^£^ p< 0.001, ^£^ ^£^ ^£^ ^£^p< 0.0001 compared to untreated h-MDM (-ML385).

To validate in human that in the early stages of infection Nrf2 activation in macrophages is dependent on ROS and in the later stages on PGE2 production, we evaluated the Nrf2 nuclear translocation in h-MDM pre-treated with GSK2795039 (NOX2 inhibitor) or with CAY10526 (PGES inhibitor) associated with AH6808 (EP2 antagonist) (Fig. 6g). Nrf2 translocation was reduced by GSK2795039 only at 4h post-*L.i* challenge and by CAY10526/AH6808 only at 48h post-*L.i* challenge (Fig. 6g). Furthermore, *L.i* proliferation was increased in h-MDM by GSK2795039 and PGE2 complementation. However, the inhibition of PGE2 production by CAY10526/ AH6808 reduced *L.i.* proliferation (Fig. 6h), demonstrating the contribution of ROS- and PGE2-mediated Nrf2 activation in the control of *L.i.* proliferation by h-MDM.

Interestingly, the pharmacological inhibition of Nrf2 by ML385 abrogated *L.i.* proliferation in h-MDM (Fig. 6i). Consistent with the anti-proliferative effect of ML385, *L.i.* lipid peroxidation, 8-isoprostane, ROS and NO production were strongly increased in infected h-MDM treated by ML385 (Fig. 6j-M).

Altogether these data validate in human that ROS/PGE2 pathway mediates Nrf2 activation and hence promotes *L.i.* proliferation through the inhibition of oxidative stress.

## Discussion

*Leishmania* parasites have developed clever strategies to resist various microbicidal mechanisms, including the oxidative burst generated by the host macrophages to limit parasite growth, upon entering host cells. We demonstrated here an increased ROS production by murine and human macrophages in the early stages of *Leishmania infantum* infection. The use of a pharmacological inhibitor of the NADPH oxidase (NOX2) reduced the Nrf2 nuclear translocation in murine and human macrophages, strengthening the contribution of ROS in early Nrf2 activation. Consistently, *L. guyanensis* was recently shown to mediate the activation of NADPH oxidase leading to trigger the release of Nrf2 from its negative regulator KEAP1 and to the translocation of Nrf2 into the nucleus (Reverte *et al*, 2021). However, during *L. infantum* infection, we showed a sustained activation of Nrf2 while the production of ROS gradually decreased over time. This indicates that in the later stages of infection, an alternative signaling is involved in Nrf2 activation. Interestingly, and in agreement with the data of Saha et *al*., (Saha *et al*, 2014) we demonstrated that in the late stages of infection, *L. infantum* enhances COX-2 and PGES gene expression in murine and human macrophages, which in turn increases the production of PGE2. In the delayed phases of *L. infantum* infection, pharmacological inhibition of PGE2 synthase and the EP2 receptor drastically reduced Nrf2 nuclear translocation in murine and human macrophages, strengthening COX-2/PGEs/PGE2/EP2r axis in the maintaining Nrf2 activation. The increase in parasite load in mice treated with the inhibitor of 15-PGDH, a prostaglandin-degrading enzyme (Huang *et al*, 2023), reinforces the importance of this signaling pathway and the crucial contribution of PGE2 *via* EP2r and not its metabolites in the progression of infection. Collectively, our data show the contribution of NOX2/ROS axis in the early activation of Nrf2 and the subsequent involvement of PGE2/EP2r in the sustainment of this activation upon *L. infantum* infection. These data were also supported in a murine model of *L. infantum* visceral infection where we demonstrated a high concentration of urinary 8-isoprostane, which reflects the oxidative stress, in the early stages post-infection, while the PGE2 metabolite levels in urine only increased in the later stages.

Genetic deletion of Nrf2 in *L. infantum*-infected mice identified this transcription factor as a central factor in the progression of visceral leishmaniasis, particularly through its role in controlling inflammation and oxidative balance. Indeed, parasite loads in the liver and spleen were significantly reduced in Nrf2^-/-^ infected mice. *L. infantum* clearance was related to a pro-inflammatory and pro-oxidant phenotype of macrophages, demonstrating that Nrf2 oriented macrophages towards an anti-inflammatory and anti-oxidant profile that promoted the progression of *L.i*. infection. Consistent with these data, besides the anti-oxidant role of Nrf2^5^, more recent data attribute an important role for Nrf2 in the suppression of macrophage inflammatory response by blocking proinflammatory cytokine transcription (Kobayashi *et al*, 2016). The adoptive transfer of Nrf2^-/-^ macrophages into infected Nrf2^+/+^ recipient mice ameliorated host response to infection, interfering that Nrf2-induced anti-inflammatory and anti-oxidant macrophages might be the cause of the systemic vulnerability of *L. infantum* infection.

Similarly, to the data obtained with Nrf2*^-/-^* mice, pharmacological inhibition of Nrf2 by the administration of ML385 in infected mice reduced the parasite load in the liver and spleen and oriented the phenotype of macrophages toward a microbicidal pro-inflammatory and pro-oxidant profile. Interestingly, the evaluation of the effect of ML385 on the infection course compared to the reference treatment Ambisome (Sundar & Chakravarty, 2010) showed a similar effect of the two molecules, highlighting the potential of ML385 for the treatment of visceral leishmaniasis.

Currently, the cell death programs of *Leishmania* genus remain poorly defined. Only a few data suggest apoptosis as a potential type of death in *Leishmania spp*. However, key apoptotic proteins involved in mammalian apoptosis have not been reported in the *Leishmania* genus (Basmaciyan & Casanova, 2019). In this work, we show that ROS-induced lipid peroxidation leads to a decrease in *L. infantum* viability, suggesting that ferroptosis could be one type of death present in *Leishmania*. To this regard, it was recently demonstrated in *Trypanosoma brucei,* a parasite belonging to *Trypanosomatidae* family as *Leishmania* species, that its death could also involve ferroptosis upon lipid peroxide accumulation (Bogacz & Krauth-Siegel). To further support that the *L. infantum* death program engaged is ferroptosis, we demonstrated that *L. infantum* infection in macrophages genetically invalidated or pharmacologically inhibited for Nrf2 induced an oxidative burst leading to lipid peroxidation of parasites and their death. Thus, the antioxidant effect of Nrf2 in macrophages protected *L. infantum* from lipid peroxidation preventing the death of parasites by ferroptosis. How macrophage Nrf2 regulates lipid peroxidation of *Leishmania* constitutes an intriguing avenue that is currently under investigation.

In conclusion, *Leishmania infantum* infection induces the successive production of ROS and PGE2 by macrophages leading to the activation of Nrf2 (Fig.7). This activation is responsible for the onset of an anti-oxidant and anti-inflammatory phenotype of macrophages favoring the progression of visceral leishmaniasis. In addition, we identify for the first time the ferroptosis-like as a cell death program of *L. infantum* and the protective effect of Nrf2 in macrophages against this parasite death. Finally, we highlight Nrf2 as a critical factor for the susceptibility of *L. infantum* infection. Synthetic inhibitor of Nrf2 activity may, hence, constitute promising compounds for the treatment of visceral leishmaniasis.

**Figure 7.**
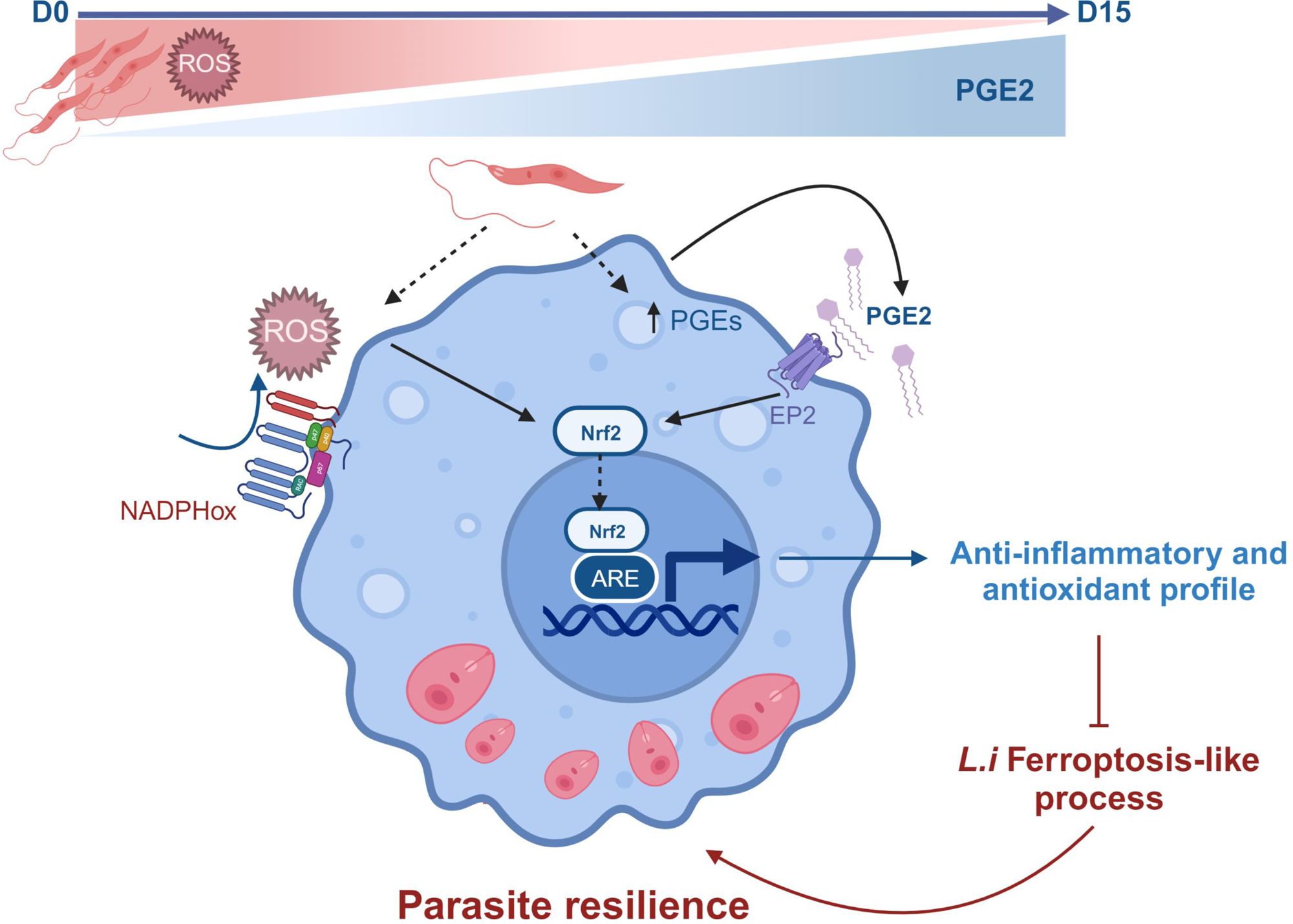
Schematic illustration of the ROS- and PGE2-mediated Nrf2 activation in macrophages leading to the *Leishmania infantum* resilience. *Leishmania infantum* infection induces the successive contribution of NOX2/ROS axis in the early Nrf2 activation and PGE2/EP2r signalling in the sustainment of Nrf2 activation. This activation is responsible for the onset of an anti-oxidant and anti-inflammatory phenotype of macrophages favoring the progression of visceral leishmaniasis. In addition, we establish macrophage-driven ferroptosis-like process as a cell death program of *L. infantum* and the protective effect of Nrf2 in macrophages against *L. infantum* killing.

## Materials and Methods

### Mice

Mice were bred and handled under protocols approved by the Conseil Scientifique du Centre de Formation et de Recherche Experimental Médico Chirurgical and the ethics board of the Midi-Pyrénées ethic committee for animal experimentation (CEEA122) with permit number 6555-2016082912056664 in accordance with European legal and institutional guidelines (2010/63/UE) for the care and use of laboratory animals. C57BL/6 mice were purchased from Janvier. Nfe2l2^+/+^ and Nfe2l2^−/−^ mice have been described earlier (Itoh *et al*, 1997).

For the *in vivo* experiments, a visceral infection was established by inoculating i.p. 50× 10^6^ stationary phase promastigotes/mouse (Lefèvre *et al*, 2013). At 14 days after infection, the liver, spleen and peritoneal macrophages were removed aseptically. The quantification of *L.i.* in tissues was done by LightCycler PCR (Roche Diagnostics) as previously described. Mice were treated with an ROS chelator (Trolox 40 mg/kg Sigma-Aldrich) every 2 days or a cocktail of PGEs inhibitor (CAY10526 5 mg/kg VWR) and prostanoid receptor antagonist 2 (AH6809 5mg/kg Sigma-Aldrich) every 2 days or with a 15-PGDH inhibitor (SW033291 at 10mg/kg Sigma-Aldrich) every 2 days, or treated with an Nrf2 inhibitor (ML385 at 10 mg/kg Sigma-Aldrich) every two days or with liposomal amphotericin B (Ambisome 3mg/kg Sigma-Aldrich) at day D0, D1, D2, D3, D4 and D10. Mice were treated with ferrostatin-1 (10mg/kg Sigma-Aldrich) each day.

For adoptive transfers, peritoneal macrophages from three Nfe2l2^−/−^ donor mice were collected and transferred (i.p.) into the corresponding WT recipient mouse model (n = 6) 10 hr before *L.i.* infection. Liver and spleens were isolated and peritoneal macrophages were harvested 14 days after infection.

### Leishmania cell culture

The cloned line of *L.infantum* (*L.i*.) (MHOM/MA/67/ITMAP-263) and the axenic amastigotes expressing luciferase activity (*L.i*-luc) have been described earlier (Lefèvre *et al*, 2013). For animal and macrophage infection, the parasites were transformed in promastigotes by changes in culture conditions (25°C and RPMI 10% FCS).

### Mouse peritoneal macrophages isolation

Resident peritoneal cells were harvested by washing the peritoneal cavity with sterile 0,9% NaCl. Collected cells were centrifuged and the cell pellet was suspended in DMEM with 5% FBS (Invitrogen). Cells were allowed to adhere for 2 h at 37°C, 5% CO2. Non-adherent cells were then removed by washing with PBS.

### Purification and generation of monocyte-derived Macrophages

Monocytes were isolated from blood Peripheral Blood Mononuclear Cells (PBMCs) from healthy donors obtained from the EFS Toulouse Purpan (France). Briefly, PBMCs were isolated by centrifugation using standard Ficoll-Paque density (GE Healthcare) according to the manufacturer’s instructions. The cells were resuspended in RPMI-1640 supplemented with 10% of foetal calf serum (FCS), 1% penicillin (100 IU/mL) and streptomycin (100 μg/ml). Monocytes were separated by negative selection using Pan monocyte isolation kit (Miltenyi Biotec). Cells were cultured in RPMI-1640 (GIBCO) supplemented with 10% FCS (Invitrogen), 100 IU/ml penicillin, 100μg/ml streptomycin, 10 ng/ml M-CSF for 5 days at 37°C under a 5% CO2 humidified atmosphere.

### Reverse Transcription and Real-Time PCR

mRNAs were isolated using the biobasic Kit (Biobasic) using the manufacturer’s protocol. Synthesis of cDNA was performed according to the manufacturer’s recommendations (Verso Kit, Thermo). RT–qPCR was performed on a LightCycler 480 system using LightCycler SYBR Green I Master (Roche Diagnostics). The primers were designed with the software Primer 3. Glyceraldehyde-3-phosphate dehydrogenase (Gapdh) mRNA was used as the invariant control. Serially diluted samples of pooled cDNA were used as external standards in each run for the quantification. Primer sequences are listed in supplemental table 1.

### Proliferation and Phagocytosis assays

Adherent murine peritoneal macrophages were prepared as previously described (Lefèvre *et al*, 2013) and pretreated 24 hours before *L.i.* challenge (parasite-to-macrophage ratio 5:1) with GSK2795039 (Sigma, 10 μM), AH6809 (Sigma, 10 μM), SW03329 (10 μM, sigma), NS398 (10 μM, Sigma), CAY10526 (10uM, VWR) or PGE2 (1uM VWR) ML385 (10µM Sigma-Aldrich). To evaluate, phagocytosis and proliferation of *L.i.*, the macrophages were challenged with *L.i*.-luc for 30 min at 37°C (phagocytosis) and for 24 hr at 37°C (proliferation). The luciferase activity was measured with a luminometer (Envison Perkin Elmer).

### ROS and NO Production, ELISA Cytokine Titration, and EIA Lipid Quantification

The ROS production by macrophages was measured by chemiluminescence in the presence of 5-amino-2,3-dihydro-1,4-phthalazinedione (luminol) using a thermostatically (37°C) controlled for 1 hr (Envision, PerkinElmer). For nitrite release, Griess reagent was used to quantify the concentration of nitrite, as previously described (Alaeddine *et al*, 2019). The cytokine releases were evaluated by ELISA in cell supernatants or in peritoneal fluids (IL-1β, IL-12, IL-6, TNFα, TGFβ, IL-10 BD Biosciences, R&D Systems). 8 isoprostane, PGE2, PGD2 and PGEM were quantified using EIA kits as recommended by the manufacturer’s protocol (Cayman).

### Immunoblotting

Total protein lysates were extracted according to standard procedures. After protein transfer, membranes were incubated with either anti-Cox2 antibody (BD transduction lab) or anti-Pges (Invitrogen 702796) or anti-Ptger2 (Invitrogen MA5-35750) or anti-β-actine (Santa cruz sc1615). Immunoblottings were revealed using a chemiluminescent substrate ECL substrate (Biorad) and images were acquired on a ChemiDoc Imaging System (Biorad).

### Fluorescence imaging confocal microscopy

Murine or human macrophages were fixed with PBS containing 4% paraformaldehyde. After permeabilization and blocking, cells were incubated at 4°C with anti-Nrf2 antibody (thermo fisher scientific MA5-42371) and then an anti-rabbit alexa 488 antibody was used (Invitrogen). Nuclei were stained with DAPI. All microscopy imagery was performed with an Operetta high content quantitative confocal imaging (PerkinElmer). For each condition, 60 000 cells were analyzed. The staining was representative of three independent experiments. At least sixty-fields/well with a minimum of 3Z planes were analyzed with integrated Columbus image analysis software. Image analysis was performed and the cells-total number, mean/well cell–Alexa Fluor intensity was computed. Nuclear translocation of Nrf2 was obtained by Alexa 488 fluorescence intensity in DAPI fluorescence intensity.

### Nuclear protein extraction and DNA-binding activity

Peritoneal macrophages from wild-type mice were previously treated for 24 hours with the NOX2 inhibitor GSK2795039 at 10µm (Sigma-Aldrich), or the PGEs inhibitor CAY10526 at 10uM (VWR) and the EP2 receptor inhibitor AH6808 at 10µM (Sigma-Aldrich). After washing with PBS-/-, macrophages were infected with a 1/5 ratio and incubated for 4h or 48h. Nuclear proteins were isolated according to the manufacturer instruction (Active Motif nuclear extraction). Nrf2 TransAM ELISA-kit (Active Motif) was used to evaluate Nrf2 DNA-binding activity. The absorbance was measured at 450nm on a microplate reader (Envision PerkinElmer).

### Leishmania Lipid peroxidation

To measure lipid peroxidation of *Leishmania infantum* parasites in macrophages, *L.i* were incubated with C11-BODIPY (581/591) (ThermoFisher) at 2 μM, in Opti-MEM medium for 2 hours at 25°C. After washes with PBS, *L.i* were resuspended in RPMI 10% FBS in the presence or absence of H_2_0_2_ at 100µM (Invitrogen) or ferrostatin-1 (Fe1) at 20µm (Sigma-Aldrich) for 4h at 37°C. Lipid peroxidation was assessed by measuring the fluorescence of C11-BODIPY at 488nm using PerkinElmer Envision plate reader.

*L.i* (pre-incubated with C11-BODIPY) were also challenged with mouse peritoneal macrophages (Nrf2 ^+/+^) or (Nrf2 ^-/-^) pre-treated or not with ML385 at 10µM (Sigma-Aldich) or h-MDM pre-treated with ML385 at 10µM (Sigma-Aldich). After 1h30 of infection, cells were washed with PBS to remove extracellular *L.i* and incubated for 4h. Lipid peroxidation was assessed by measuring the change in fluorescence of C11-BODIPY at 488nm using direct fluorescence reading with the plate reader (Envision PerkinElmer).

### Cell viability

Cell viability of *L.i*-infected macrophages or *L.i* promastigotes alone was assessed using the AlamarBlue® proliferation assay (Invitrogen) according to the manufacturer’s instructions. AlamarBlue reagent was incubated for 4 hours. The fluorescence intensity was measured with a fluorescence plate reader (Varioskan Thermo fisher scientific) with an excitation of 560 nm and an emission of 590 nm. The results were expressed as the percentage of viable cells compared to uninfected macrophages.

### Statistical analysis

Statistical analysis was performed using the means multiple comparison method of Bonferroni-Dunnett test. P <0.05 was considered statistically significant. Data were expressed as mean ± SEM.

## Acknowledgements

We thank Philippe Batigne (Université Paul Sabatier) for excellent technical support and Jean Loup Lemesre (UMR INTERTRYP, Montpellier) for providing the cloned line of *L.infantu*m (MHOM/MA/67/ITMAP-263) expressing luciferase activity.

## Author contributions

**Clément Blot**: Conceptualization; formal analysis; investigation; methodology; writing–original draft; writing–review and editing. **Kimberley Coulson**: Formal analysis; investigation. **Marie Salon**: Investigation; methodology. **Margot Tertrais**: Investigation; methodology. **Rémi Planès**: Investigation; methodology. **Karin Santoni**: Investigation; methodology. **Hélène Authier**: Investigation. **Godefroy Jacquemin**: Investigation. **Mouna Rahabi**: Investigation. **Mélissa Parny**: Methodology. **Isabelle Raymond Letron**: Methodology; formal analysis. **Etienne Meunier**: Conceptualization; supervision; writing–original draft; writing–review and editing. **Lise Lefèvre**: Conceptualization; supervision; validation; writing–original draft; writing–review and editing. **Agnès Coste**: Conceptualization; supervision; validation; funding acquisition; project administration; writing–original draft; writing– review and editing.

## The paper explained

### Problem

*Leishmania* species are intracellular protozoan parasites that can cause severe visceral disease. *Leishmania* parasites have developed clever strategies to resist microbicidal mechanisms, including the oxidative burst generated by the host macrophages to limit parasite growth. Nuclear factor erythroid 2-related factor 2 (Nrf2), is among the key factors that regulate the cellular redox balance and previous work has showed the activation of Nrf2 by *Leishmania spp*. However, the precise molecular mechanisms responsible for activating Nrf2 in macrophages and its role in the course of the infection remains to be elucidated.

### Results

Using both genetically Nrf2-deficient mice and a pharmacological approach on murine and human macrophages, we have identified the early implication of reactive oxygen species (ROS) in NRF2 activation and the pivotal role of PGE2 eicosanoids in the maintenance of Nrf2 activation. Consequently, the macrophage phenotype is oriented toward a protective anti-oxidant and anti-inflammatory profile that prevents the death of parasites by ferroptosis-like process and thus promoting *L.i.* infection.

### Impact

In this study, we highlight Nrf2 as a critical factor for the susceptibility of *L. infantum* infection. Synthetic inhibitor of Nrf2 activity may, hence, constitute promising compounds to improve treatment of visceral leishmaniasis.

---

**Table 1:**
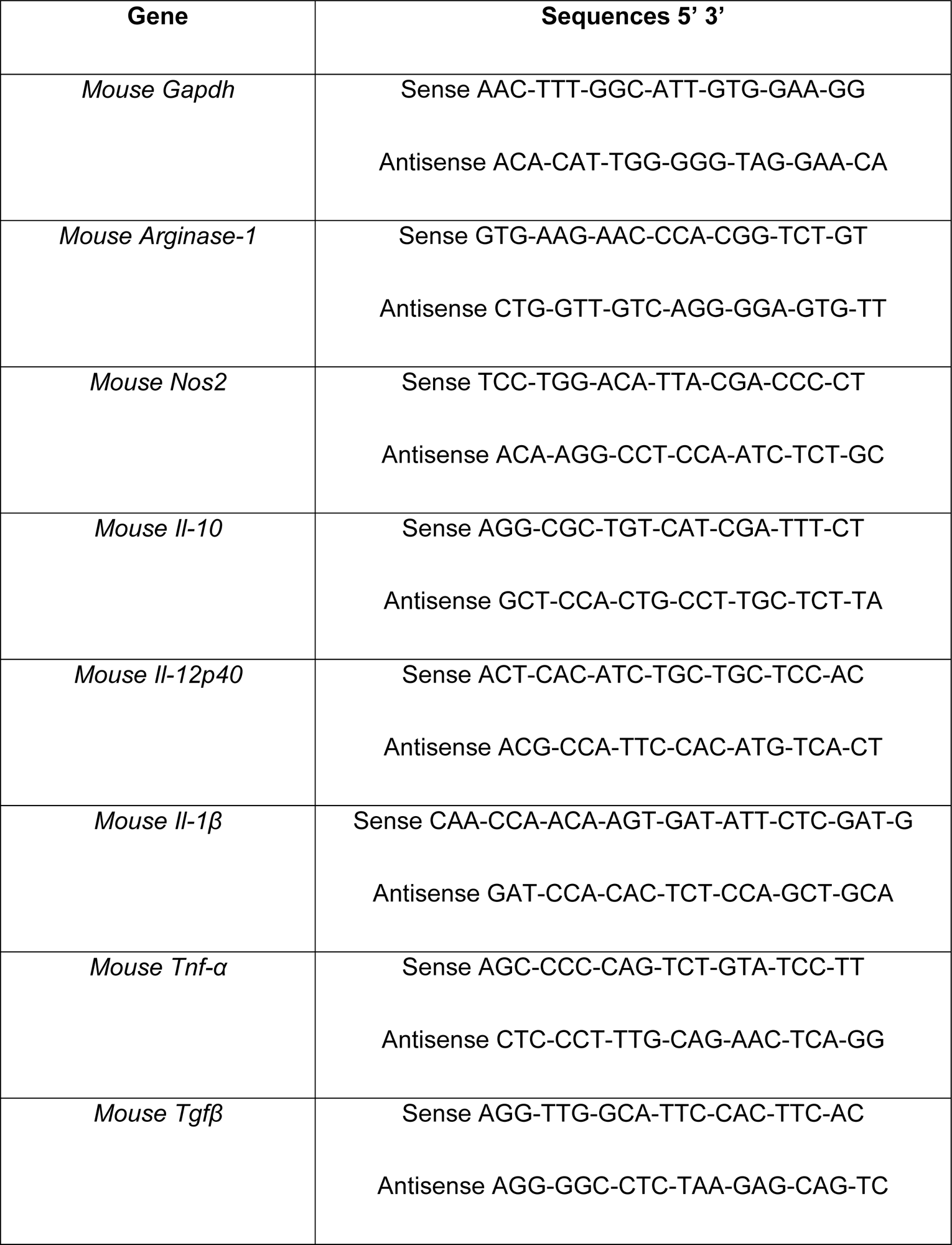
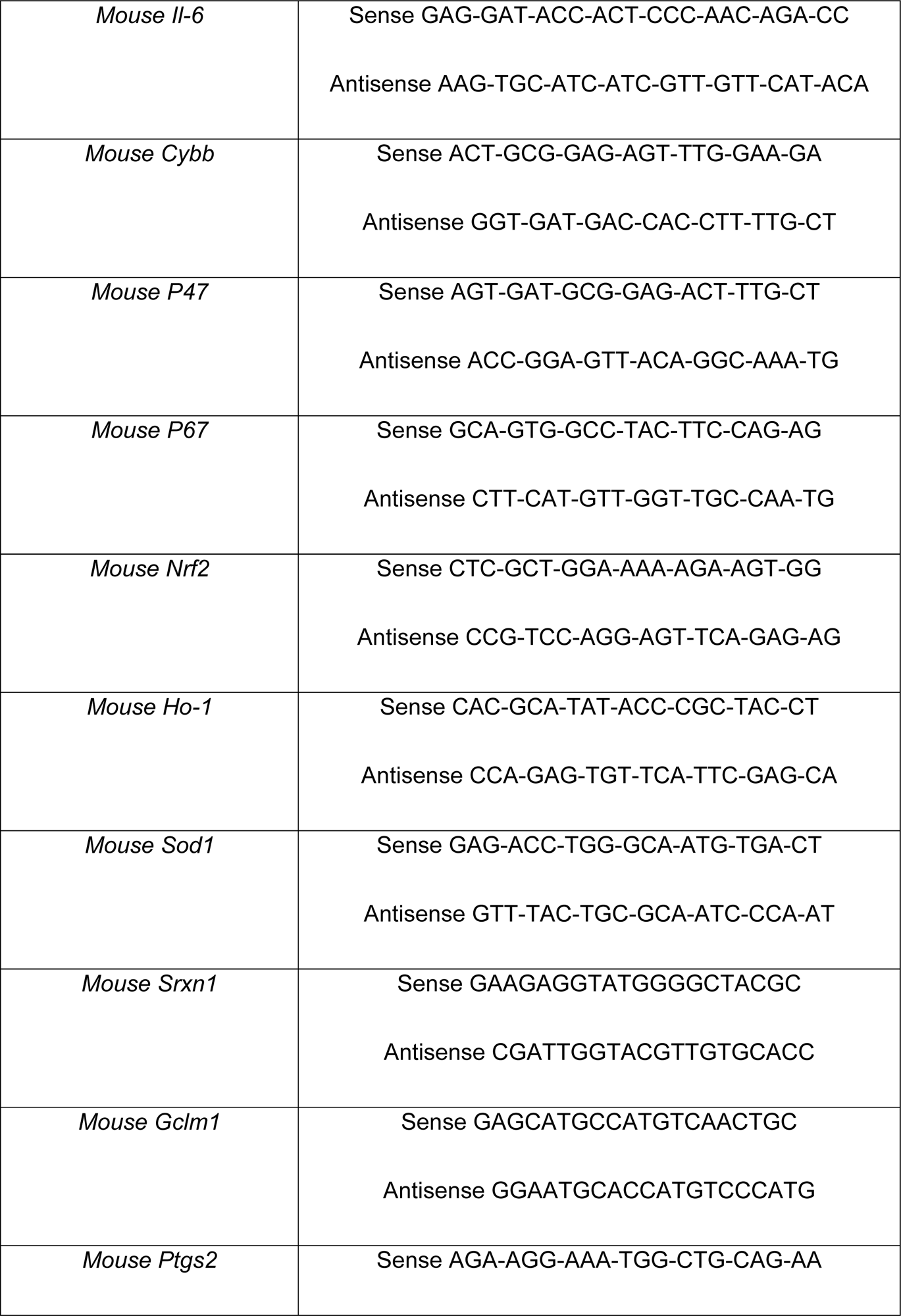
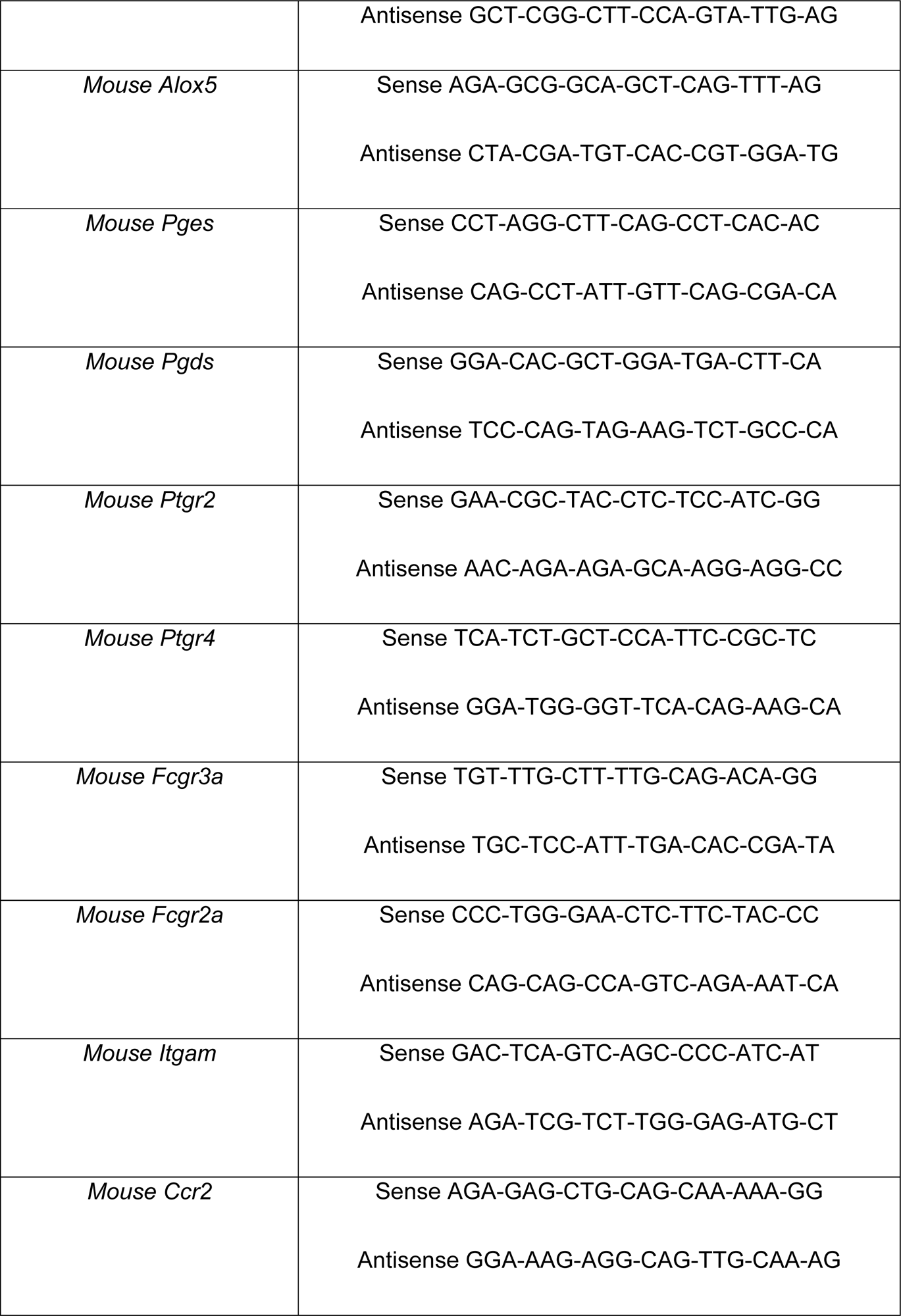
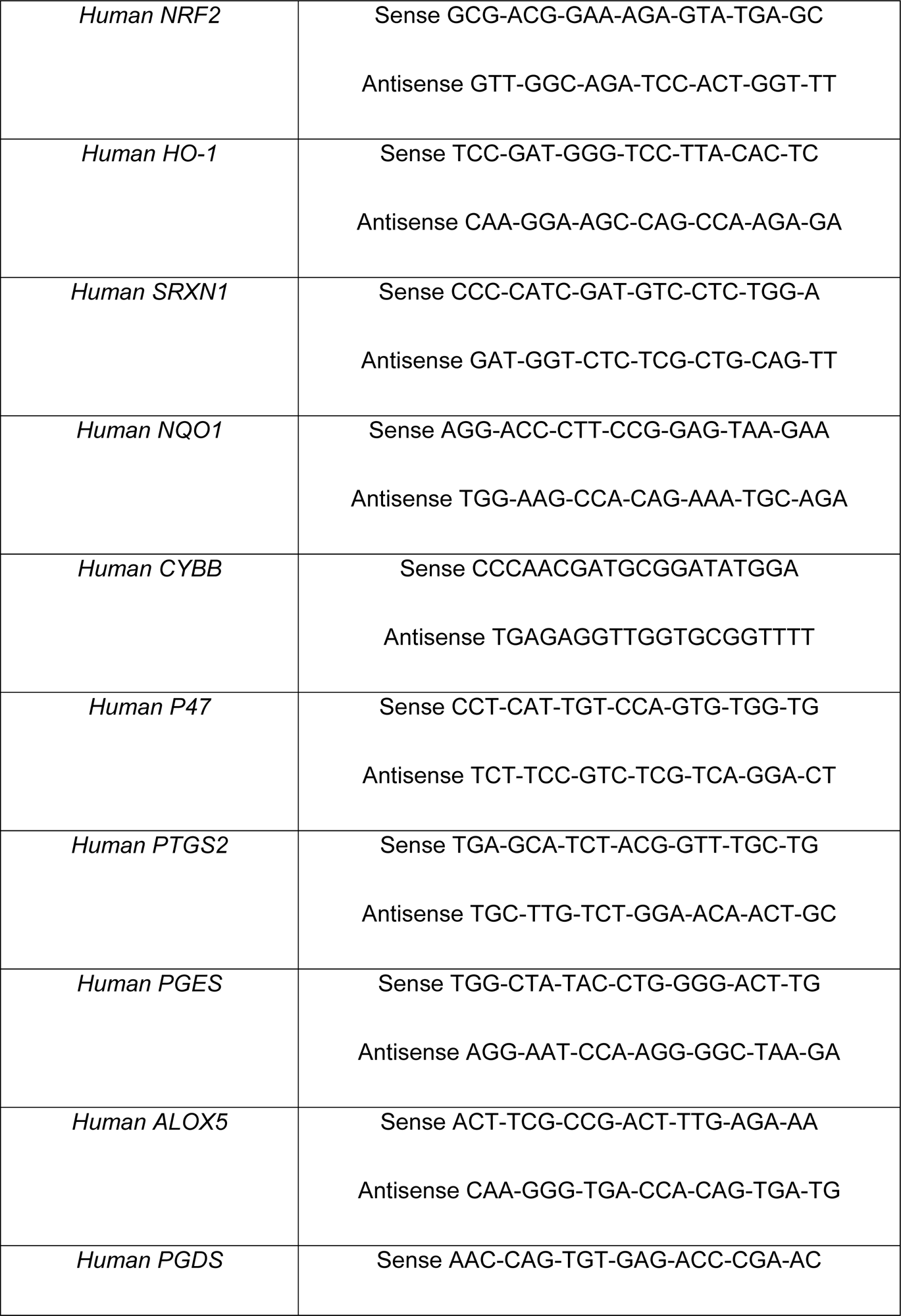
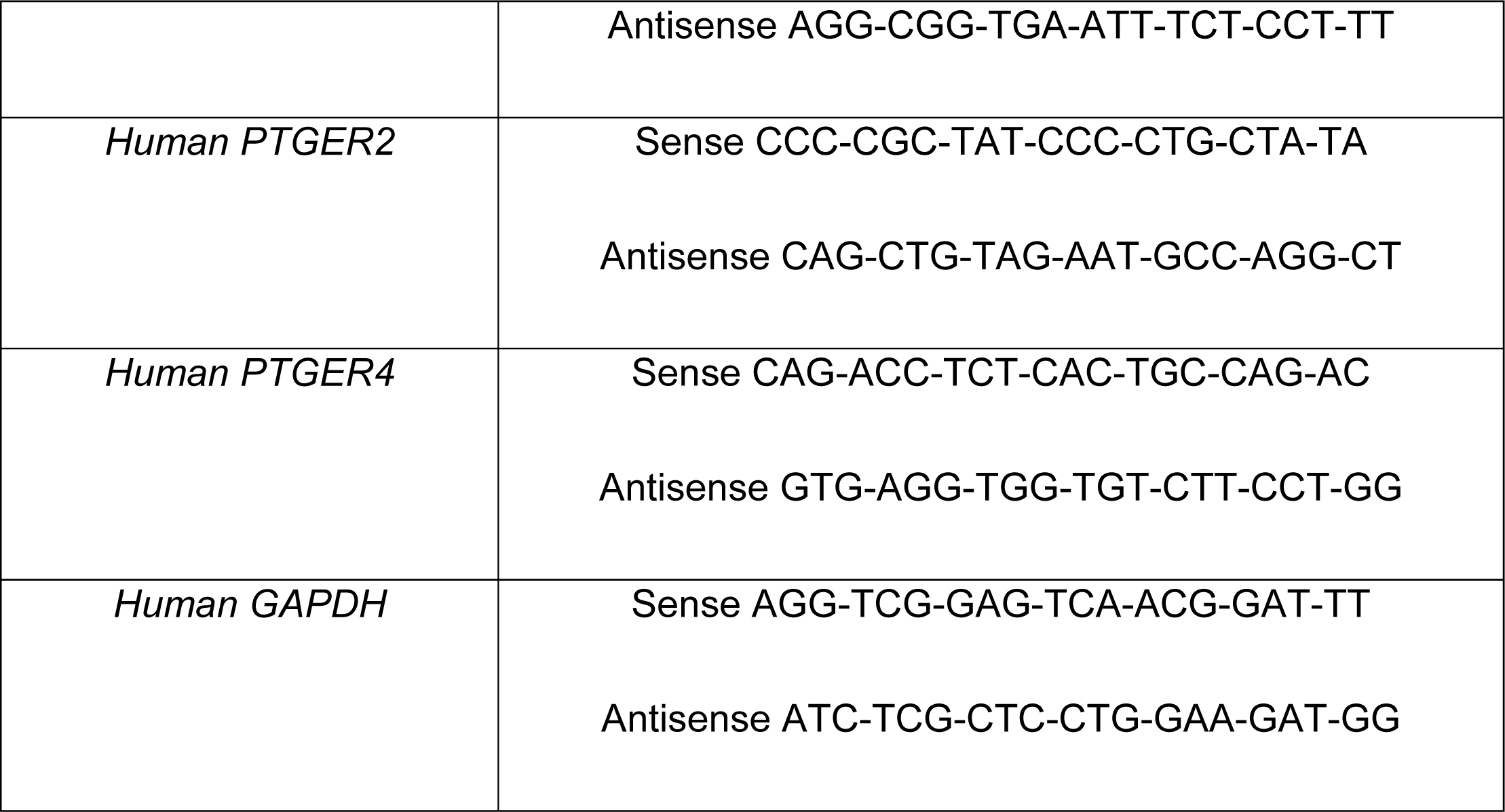
Primer sequences.

